# Transcranial magnetic stimulation during British Sign Language production reveals monitoring of discrete linguistic units in left superior parietal lobule

**DOI:** 10.1101/679340

**Authors:** David Vinson, Neil Fox, Joseph T. Devlin, Karen Emmorey, Gabriella Vigliocco

## Abstract

Successful human hand and arm movements are typically carried out by combining visual, motoric, and proprioceptive information in planning, initiation, prediction, and control. The superior parietal lobule (SPL) has been argued to play a key role in integrating visual and motoric information particularly during grasping of objects and other such tasks which prioritise visual information. However, sign language production also engages SPL even though fluent signers do not visually track their hands or fixate on target locations. Does sign language production simply rely on the motoric/ proprioceptive processes engaged in visually guided action, or do the unique characteristics of signed languages change these processes? Fifteen fluent British Sign Language users named pictures while we administered transcranial magnetic stimulation (TMS) to left SPL, a control site, or no TMS. TMS to SPL had very specific effects: an increased rate of (sign-based) phonological substitution errors for complex two-handed signs (those requiring hand contact), but TMS did not slow or otherwise impair performance. Thus, TMS decreased the likelihood of detecting or correcting phonological errors during otherwise successful bimanual coordination, but it did not noticeably alter fine movement control. These findings confirm that for fluent signers SPL has adapted to monitor motor plans for discrete hand configurations retrieved from memory as well as more fine-grained aspects of visually guided actions.

## 1. Introduction

Hand actions can be fast, accurate, highly complex, and most importantly, controlled and corrected in real time. Often, this requires taking into account information from multiple sensory modalities, particularly vision and proprioception, that may change rapidly over time. Evidence from a wide variety of studies implicates the superior parietal lobule as playing a key role in controlling human hand and arm actions, in particular by maintaining body-centric representations during actions that must be constantly updated based on visual or proprioceptive information (Jackson and Husain, 2006). For most hand actions studied, vision is essential, as reaching and grasping a potentially movable object requires tracking its position in space, along with visual information about hand position (Battaglia-Mayer et al., 2014). Proprioceptive cues are also available, which under normal circumstances may be redundant with vision, but become increasingly relevant in situations of occlusion or inconsistency (Bernier and Grafton, 2010; Parkinson et al., 2010; Tagliabue and McIntyre, 2014), or when motor plans must be altered to achieve successful action (Della-Maggiore et al., 2004; Reichenbach et al., 2014) in which cases SPL activity is increased. Most studies, however, focused only upon visually-guided hand actions, in which movement to a target location depends on accounting for the position of an external target and thus visual information is still relevant. It remains unknown whether comparable processes underlie production of hand movements generated from internal representations and which are not visually guided such as sign language or gesture production.

Some studies of reaching and grasping manipulate the availability of visual cues and/or their consistency with proprioceptive cues (e.g. Jackson and Husain, 2006; Reichenbach et al., 2014). The findings from this research suggest different action control systems (Chib et al., 2009) with proprioceptive-only systems distinct from combined sensory systems that take both types of information into account in movement correction. The tasks, however, are still explicitly visual in that specific visual properties of the target must determine the motor plan. To avoid this issue, Parkinson et al. (2010) asked blindfolded participants to make reaching hand movements to a button box near their waist, thus planning actions based on bodily locations in memory. Increased SPL activation was found when movement to a new target location was required. If SPL constantly maintains a predictive model of arm/hand position based on efferent copy of the actual motor command (Wolpert et al., 1998), this effect would arise because of the greater need to incorporate bodily position into plans/forward models when moving to a new location. However, this study involves a relatively unfamiliar action and it is unclear whether “new” and “familiar” movements exhibited comparable kinematic properties, potentially undermining the conclusion that SPL’s engagement relates to internal models, rather than properties of overt execution.

We suggest that motoric components can be investigated far more directly with a highly-practiced form of action that is not visually guided: sign language. Research comparing signed and spoken language production has revealed that despite fundamental differences between visual-manual and acoustic-oral modalities, similar left-hemisphere language networks are engaged in production of both (Braun et al., 2001; Emmorey et al., 2007; 2016). Key differences are also observed, in particular with respect to activation of left SPL in the production of sign but not speech (Braun et al, 2001; Emmorey et al, 2002; 2004; 2007; 2014; 2016; Kassubek et al., 2004; Corina and Gutierrez, 2017). Crucially, signers rarely look at their own hands (Emmorey et al., 2009a), and visual feedback is not used to monitor sign articulation (Emmorey et al., 2008; 2009b) – although hands and arms are regularly visible in the periphery, they are not the focus of visual attention. Thus, proprioception is hypothesized to dominate the process of sign articulation. Moreover, unlike reaching and grasping, the details of hand configuration and movement in signing are not determined by properties of a visual target, but instead are retrieved from memory. As is the case for spoken words, lexical signs are made up of relatively small phonological units that are not meaningful in themselves and are combined to produce meaningful content (Stokoe, 1960/2005). In terms of manual articulation, three major parameters are relevant: hand configuration (handshape and orientation), place of articulation, and movement (Sandler and Lillo-Martin, 2006). To produce a sign, these features must be retrieved on the basis of a signer’s intentions and carried out with minimal visual attention to the hands or their target location(s). If SPL engagement is due to integrating visual and proprioceptive information, why would SPL activity be seen during sign production?

Emmorey and colleagues (2004; 2007; 2014; 2016) hypothesized that increased activity in left SPL during sign production is due to self-monitoring: more specifically, generating predictions and monitoring hand position, especially relative to other locations on the body via proprioceptive feedback (see Pickering and Garrod, 2013; Hickok, 2014, for similar approaches to speech monitoring). Crucially, these predictions do not faithfully reproduce all motoric aspects of production, but incorporate simplified aspects of production specifically relevant for different levels of linguistic representation/processing such as phonemes, syntactic class, and semantics.

Consistent with the notion that SPL activity in signing reflects prediction rather than overt action, it is also found in studies employing covert naming in which signers are asked only to imagine producing a sign (Kassubek et al., 2004; Hu et al., 2011). Left SPL is engaged even in covert sign production, suggesting a role in motor imagery and not only explicit motoric behaviour (Saiote et al., 2016), consistent with SPL engagement in generating predictions/forward models for error monitoring. As increased SPL activity is also observed during sign (but not speech) comprehension (Leonard et al., 2012), these same internal predictions have been argued to be employed in comprehending others’ signs as well as guiding one’s own production (Emmorey et al., 2014), with movement simulation playing a key role. A study of BSL comprehension (Watkins and Thompson, 2017) demonstrated that signers’ own bodily representations are relevant for more complex signs. While all signers were faster to recognise one-handed signs produced by a right-handed signer, left-handers were faster to recognise non-symmetrical two-handed signs produced by someone whose handedness matched their own (but see Corina and Gutierrez, 2017). These findings from sign comprehension fit well with a PET study of sign production (Emmorey et al., 2016) showing increased in SPL activity for one-handed signs that contacted the body, compared to one-handed signs or two-handed signs produced in neutral space.

We sought to identify the processes engaged in non-visually guided action by interfering with left superior parietal lobule (SPL) activity during British Sign Language (BSL) picture naming using TMS. Picture naming is particularly useful for this purpose, as it yields not only naming latencies and accuracy, but also different errors reflecting different stages of lexical retrieval (Klima and Bellugi, 1969; Fromkin, 1971; Garrett, 1975; Hohenberger et al., 2002) offering more nuanced insight into the specific aspects of action for which SPL is engaged. While picture naming is visually initiated, actions to produce a sign are not visually guided, nor is the picture a target of the action.

If SPL is reliably engaged in retrieval of BSL signs or initiation of overt sign production, applying TMS should broadly affect both speed and accuracy: slowing response initiation relative to a control condition, and increasing the rate of naming errors. Because previous studies, both of sign language production and other types of actions, suggest greater SPL engagement during complex coordination or precise target-directed movement (Bernier and Grafton, 2010; Parkinson et al., 2010; Tagliabue and McIntyre, 2014; Emmorey et al., 2016), TMS effects on response times might be limited to signs requiring coordinated movement of both hands or precisely-defined place of articulation. However, if SPL is more engaged in generating internal plans for monitoring language-specific forms (Emmorey et al., 2004; 2007; 2014; 2016), effects on response initiation should not occur. Rather, TMS to SPL should elevate the rate of slips of the hand, with the type of error reflecting the nature of the representations/forward models that are being generated. Based on the literature on reaching and grasping (Jackson and Husain, 2006; Parkinson et al., 2010; Reichenbach et al., 2014), precision movement control is most likely to be affected, in which case TMS should increase the rate of manual dysfluencies (failure to finely correct movement trajectory or hand configuration during production) resulting in slight deviations from target signs. Alternatively, the literature on self-monitoring in speech (Pickering and Garrod, 2013; Hickok, 2014) suggests we should observe failures to detect and correct errors involving higher-level linguistic representations such as substituting discrete phonological features (producing a sign with a different hand configuration, location, or movement, as illustrated in Figure 1), or semantic substitutions (e.g. signing OWL for EAGLE).

**Figure 1.**
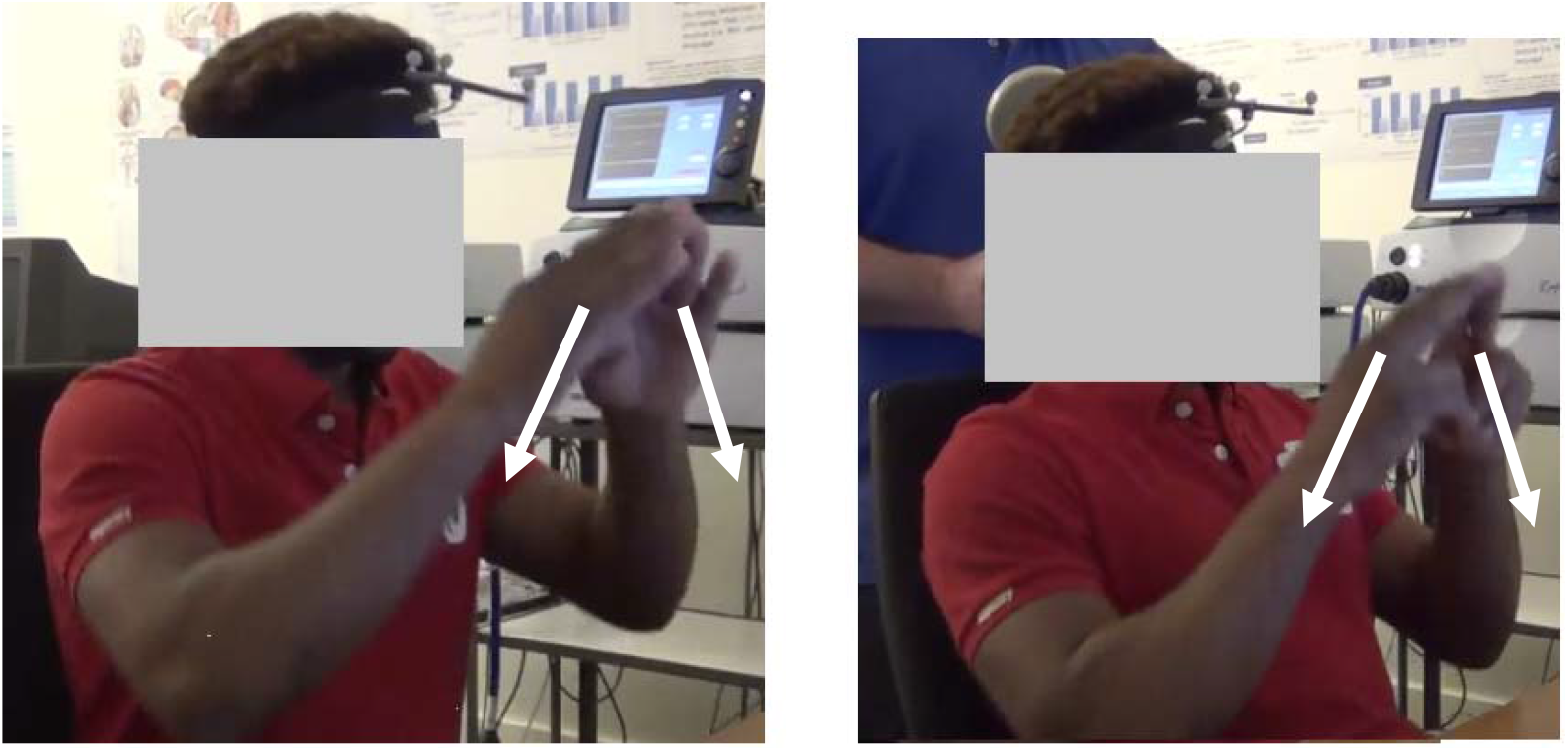
Example of a phonological feature substitution error. Left panel: BSL sign MOUNTAIN during initial name agreement phase. Right panel: BSL sign MOUNTAIN during TMS study (note coil location on left SPL, and modification of handshape from the correct utterance).

## 2. Methods

### 2.1. Participants

Fifteen fluent BSL signers (five women, 10 men; age range 19-45) participated in the study. Four were hearing, 11 deaf from birth. Most (n=12) were native signers, and all had been using BSL for more than 15 years and rated their own BSL skills as 6/7 on a 1-7 scale. The hearing signers were all employed as BSL-English interpreters at the time of testing. The study was approved by UCL Research Ethics committee; all participants gave their informed consent to participate.

### 2.2. fMRI study of covert naming

A functional MRI study was first carried out to localise each individual’s peak of activity in SPL during covert sign production. Imaging was conducted on a Siemens 1.5 Tesla MR scanner at the Birkbeck-UCL Centre for Neuroimaging (BUCNI) in London. The functional data were acquired with a gradient-echo echo planar imaging (EPI) sequence (TR = 3000 ms; TE = 50 ms, FOV = 192 × 192, matrix = 64 × 64, 35 slices), resolution of 3 × 3 × 3 mm: one run of 80 volumes (4 min). Two additional functional runs were subsequently obtained for a different study. A high-resolution anatomical scan was also obtained: (T1-weighted FLASH, TR = 12 ms; TE = 5.6 ms; 1 mm^3^ resolution).

Functional data were obtained for covert picture naming: participants were presented with a series of line drawings (Snodgrass and Vanderwart, 1980) and were asked to imagine producing the picture names in BSL without moving their hands or bodies. Each picture (n=40) appeared for 500msec, followed by a blank interval (jittered intervals, mean: 3381msec, range: 2000-4905). After each block of 10 trials a rest block of 15sec occurred.

Data processing was carried out using FSL 4.0 (Woolrich et al., 2009). Data were realigned to remove small head movements (Jenkinson et al., 2002) and smoothed (Gaussian kernel FWHM = 6 mm) before entering the images into a general linear model with two conditions of interest: covert naming and rest. Blocks were convolved with a canonical hemodynamic response function (Glover, 1999). Estimated motion parameters were also entered as covariates of no interest to reduce systematic variance related to head motion. The linear contrast covert naming > rest was the measure of interest. In each participant we identified the peak of activity for covert naming in left SPL by taking the highest Z-valued voxel within an individual’s left SPL for the covert naming > rest comparison. This voxel would serve as the target for TMS to left SPL in the subsequent session. Mean coordinates were MNI (−21, −67, 58); standard deviations (6.1, 7.8, 7.6mm), coordinates for individuals appear as Supplemental Table 1.

### 2.3. TMS experiment

The TMS experiment was conducted at least one week after the fMRI study for each participant. Materials for the study were 204 line drawings of objects (Snodgrass and Vanderwart, 1980 or created in similar style): items for which BSL lexical signs exist, in most cases using materials that had been normed in the process of other BSL studies (Vinson et al., 2008; 2010; 2015) and which varied widely across phonological characteristics such as hand configuration, place of articulation, and movement. We avoided signs that are produced on the upper part of the head or face to avoid interfering with the TMS configuration, compound signs, and any items expected to elicit English fingerspellings instead of lexical signs. An additional 12 pictures were used for practice.

The study began with a name agreement phase (no TMS). Each picture was presented one at a time, and participants were asked to name each picture using their preferred BSL sign. Presentation was self-paced, using the same display configuration that would also be used in the upcoming TMS study. The participant pressed keys on the keyboard with both hands and held them down until ready to produce a sign. This allowed us to familiarise participants with the pictures, as well as identifying each individual’s preferred sign for a given picture -- essential as there is substantial variation in BSL lexical forms (Stamp et al., 2015). For the main experiment, key release latency served as a measure of sign production onset. Pictures were presented in a random order using E-Prime 2.0 (Schneider et al., 2012). Once this initial naming phase was complete, we proceeded to the TMS phase: picture naming with the TMS coil in place. We first demonstrated the TMS equipment, including sample stimulation to the target site, to ensure the participant was comfortable with stimulation and wished to continue with the main study.

The crucial question concerns the effects of TMS to left SPL during BSL production. We compared this to the effects of TMS at a control site (vertex), as well as naming without TMS: 2×2 design with independent variables location of TMS coil (left SPL, vertex) and TMS (present, absent). On TMS-present trials 4 pulses of TMS was administered at 10 Hz: the first at picture onset and the next three at 100, 200 and 300ms post-picture-onset.

Left SPL was targeted on the basis of each individual’s fMRI activation peak (covert naming > rest); the vertex was targeted on the basis of anatomical landmarks. Stimulation used a Magstim Rapid^2^ stimulator (Magstim, Whitland, UK) with a 70mm figure-of-eight coil. Accurate targeting was achieved through use of a frameless stereotaxy system (Brainsight software) and a Polaris Vicra infrared camera (Northern Digital, Waterloo, Ontario, Canada).

TMS without picture naming was carried out first to acclimate participants and ensure they wished to continue. Twelve practice trials then familiarised the participant with the experience of TMS during naming, using the same display characteristics as the main experiment. TMS intensity was determined on the basis of pilot studies and was constant across individuals at 65% of maximum stimulator output. None of the participants reported any discomfort from stimulation at either left SPL or vertex, and all wore ear plugs throughout the TMS study.

The experiment was carried out using a blocked design: there were four blocks, each with a fixed location of the TMS coil (rotated between participants in an ABBA design; e.g. targeting left SPL on blocks 1 and 4, and vertex on blocks 2 and 3 for the first participant, and vice versa for the second). Within each block, TMS was pseudorandomly applied, such that there were an equal number of TMS and no-TMS trials per block, and never more than three trials with TMS in a row. A ten-minute break occurred between the second and third blocks.

The experimenter pressed a foot pedal to begin each block, thus ensuring the coil was correctly in place before the trials began. The foot pedal was also available for the experimenter to immediately interrupt the study (for example, to reposition the coil if needed, or to immediately stop in case of potential participant distress). The experimenter held the coil in place and monitored its positioning via an active targeting display using Brainsight on a screen visible to the experimenter but occluded from the participant’s view by the stimulus display screen. A second experimenter monitored the participant and communicated with the participant and the other experimenter using BSL. Each trial was initiated by the participant pressing and holding keys on the computer keyboard with both hands. Once a key was pressed a fixation screen appeared for 400msec, followed by the picture for that trial which remained on screen until key release was detected. On TMS trials, onset of the first TMS pulse was synchronised to picture onset. Once a key release was detected, the screen remained blank for 1500msec, then a fixation screen appeared as a signal that the next trial was ready (see Figure 2).

**Figure 2.**
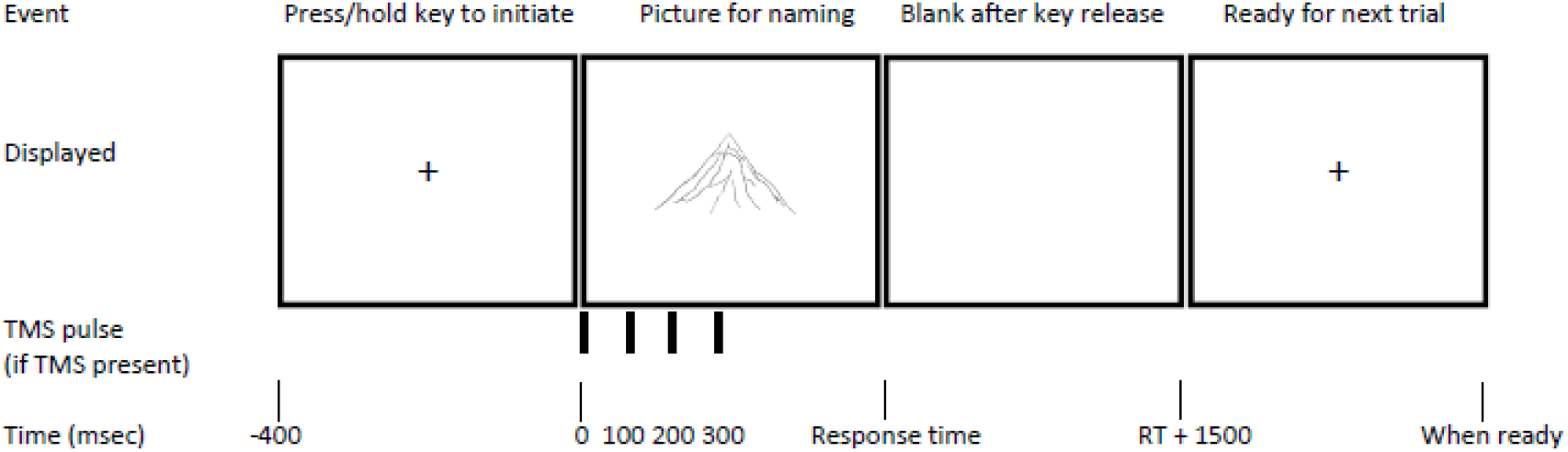
Illustration of the sequence and timing of events on a trial (with TMS present).

Position/trajectory of TMS was recorded on each TMS trial using Brainsight: the signal from the controller PC to initiate TMS also logged the coil position, stimulation trajectory and distance to the target site. All trials were video-recorded using two cameras, one on each side of the participant, approximately 45° from the participant’s eyes to centre of screen, and analysed later for accuracy.

### 2.4. Design and analysis

Errors and dysfluencies were analysed for all participants and were classified into three types (Vinson et al., 2010). Coding of errors was done blind to the presence/absence of TMS (but not coil location, as this was visible in the video recordings) and included the following classifications. <u>Lexical substitution errors</u> were cases in which a participant produced an identifiable lexical sign that was not the target (and which did not match their productions in the initial naming phase). Nearly all of these were immediately self-corrected, often before completion of the erroneous sign. <u>Phonological feature errors</u> were productions in which the resulting utterance was not a lexical sign (verified against the participant’s production in the initial naming phase, and in some cases further verified by follow-up checks with the individual), and also included clear substitution of a phonological element (e.g. hand configuration, place of articulation, movement). Sixty percent of these involved an incorrect hand configuration (handshape/orientation), 32% incorrect movement, and 8% incorrect place of articulation. For an example see Figure 1. <u>Dysfluencies & hesitations</u> were any other noticeable errors or problems not counted as lexical substitutions or phonological feature errors. This included cases of visible pauses after key release but before sign production, as well as other inaccurate or nonstandard productions that did not involve substitution of phonological features. This category also included one-handed signs for which both hands were lifted, and two-handed signs with delayed lifting of the second hand.

Correct naming latencies and overall accuracy were analysed using 2×2 ANOVA (stimulation location × presence of TMS). To test for the comparable interaction on different types of errors, we employed non-parametric tests (Wilcoxon signed-rank test) on difference scores (lSPL: TMS - no TMS, vs. Vertex: TMS - no TMS) per error type, necessary because the occurrence of errors deviates highly from normality. All analyses were conducted with participants as random effect (indicated by subscript 1) and items as random effect (subscript 2). For errors we employed directional tests under the assumption that stimulation to SPL should increase error rate if it is engaged in ensuring correct production, tested against an alpha level of .05. Analyses were carried out using IBM SPSS 20.

## 3. Results

### 3.1. Preliminary data processing

We excluded 16 trials (0.5%) on which a participant did not produce a sign - typically due to early release of the keyboard. For analysis of naming latencies (measured as the time at which the first key was released) we excluded one participant whose responses were extremely deliberate (mean RT > 2000ms vs all others mean RT < 1000ms). We also excluded errors or dysflencies (8.9% of all trials) from analysis of response latencies as well as excessively fast or slow trials: faster than 300msec or slower than 5000msec (0.7%).

### 3.2. Naming latencies and errors

Analysis of correct naming latencies revealed no effects of TMS: main effect of coil location (F_1_, F_2_ < 1); Main effect of TMS (F_1_(1,13) = 4.253, p=.060; F_2_<1); Interaction F1(1,13) = 1.023, p=.330; F2<1). Responses were highly accurate overall (91.1%). Errors were classified into three types: lexical substitution errors (e.g. signing OWL instead of EAGLE: 2.3% of all trials), phonological feature errors (producing a sign with a different hand configuration, location, or movement: 3.7%), or dysfluencies/ hesitations (2.1%). A 2×2 ANOVA of overall error rate (combining all error types) revealed no general effects of TMS on overall accuracy (all F <1). Looking separately at the different error types, there was a difference in the effects of TMS to left SPL (compared to vertex) but only for phonological feature errors. For lexical errors and dysfluencies there was no reliable interaction between coil location and presence of TMS (subjects and items |Z|<1 in both cases). However, for phonological feature errors there was a reliable interaction (Z_1_=2.60, p=.004; Z_2_=1.76, p=.039) indicating that TMS to SPL selectively increased the number of phonological feature errors while leaving the other two error types unaffected; see Figure 3 for overall response times and errors of different types by experimental condition. Because previous literature implies increased engagement of SPL in producing signs requiring coordination of hand/s with body locations (Emmorey et al., 2016; Parkinson et al., 2010; Striemer et al., 2011), we divided the signs into those produced with one or two hands, and those that involved hand-to-hand contact or no contact (there were insufficient numbers of signs produced with hand-to-body contact). Signs without contact showed no effects of TMS (both |Z|<1), but the number of phonological feature errors more than doubled under TMS to left SPL for two-handed signs with hand-to-hand contact (Z_1_=1.90, p=.023; Z_2_=1.98, p=.024); see Table 1.

**Figure 2.**
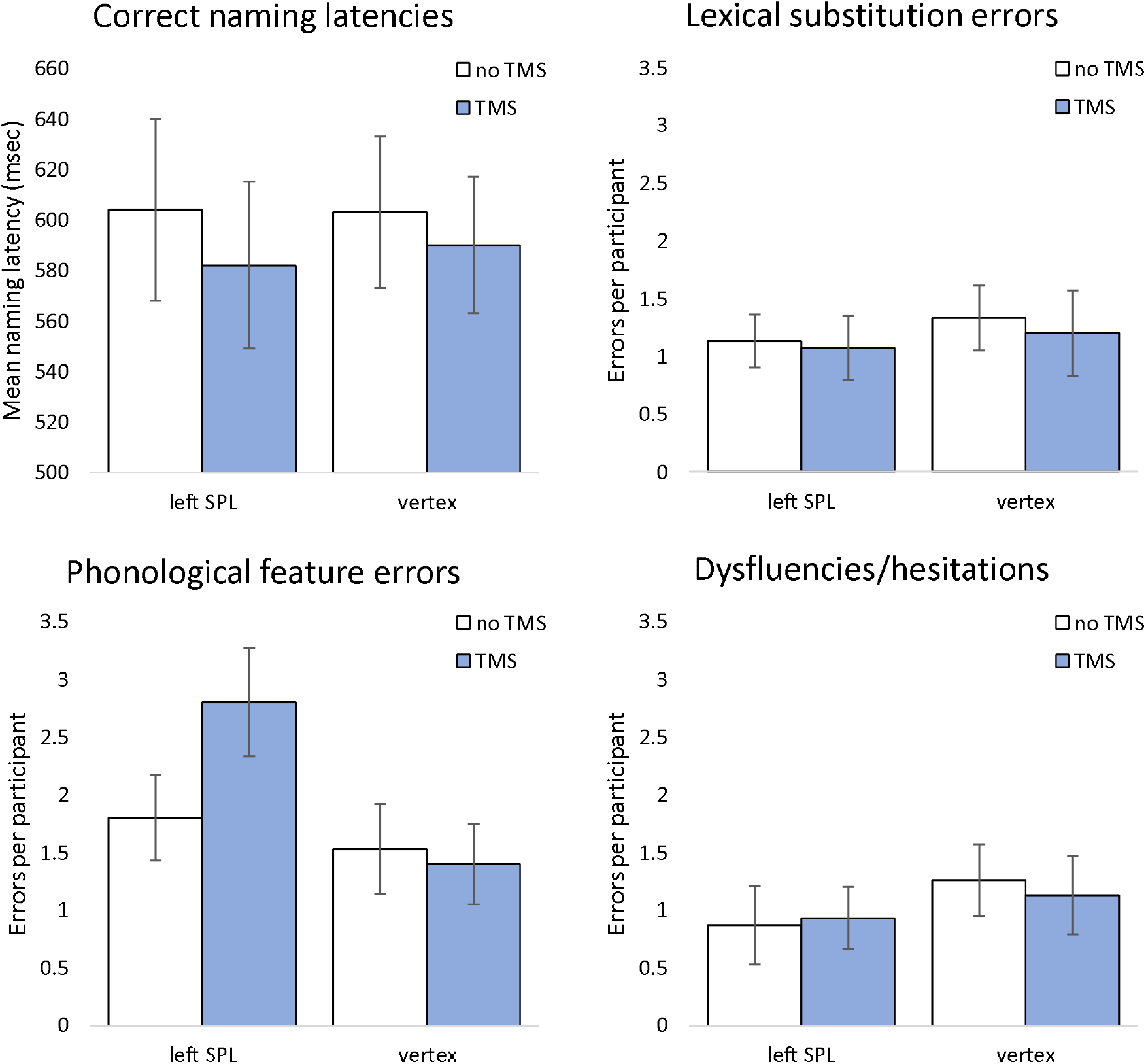
Effects of coil location and presence of TMS on naming latencies and different error types. Error bars: standard error of mean based on analysis by participants. Upper left panel: naming latencies. Upper right: lexical substitution errors. Lower left: phonological feature errors. Lower right: dysfluencies/hesitations. TMS: transcranial magnetic stimulation. SPL: superior parietal lobule.

**Table 1.** Number of errors of different types broken down by hands (one-handed or two-handed) and type of contact (body contact, hand contact, or none), as a function of TMS and coil location. Upper panel: phonological feature errors. Middle panel: lexical-semantic errors. Lower panel: dysfluencies. TMS: transcranial magnetic stimulation. SPL: superior parietal lobule.

Phonological errors
|  | One-handed signs |  |  |  | Two-handed signs |  |  |  |  |  |
| --- | --- | --- | --- | --- | --- | --- | --- | --- | --- | --- |
|  | Body contact |  | No contact |  | Body contact |  | Hand contact |  | No contact |  |
|  | No TMS | TMS | No TMS | TMS | No TMS | TMS | No TMS | TMS | No TMS | TMS |
| SPL | 3 | 3 | 9 | 13 | 1 | 3 | 7 | 20 | 7 | 3 |
| Vertex | 1 | 1 | 8 | 5 | 0 | 2 | 7 | 7 | 7 | 6 |

Lexical-semantic errors
|  | One-handed signs |  |  |  | Two-handed signs |  |  |  |  |  |
| --- | --- | --- | --- | --- | --- | --- | --- | --- | --- | --- |
|  | Body contact |  | No contact |  | Body contact |  | Hand contact |  | No contact |  |
|  | No TMS | TMS | No TMS | TMS | No TMS | TMS | No TMS | TMS | No TMS | TMS |
| SPL | 1 | 3 | 5 | 3 | 1 | 0 | 6 | 5 | 4 | 4 |
| Vertex | 3 | 1 | 4 | 6 | 0 | 0 | 8 | 7 | 5 | 4 |

Dysfluencies
|  | One-handed signs |  |  |  | Two-handed signs |  |  |  |  |  |
| --- | --- | --- | --- | --- | --- | --- | --- | --- | --- | --- |
|  | Body contact |  | No contact |  | Body contact |  | Hand contact |  | No contact |  |
|  | No TMS | TMS | No TMS | TMS | No TMS | TMS | No TMS | TMS | No TMS | TMS |
| SPL | 2 | 0 | 3 | 6 | 0 | 1 | 4 | 4 | 4 | 3 |
| Vertex | 1 | 1 | 3 | 5 | 0 | 1 | 9 | 5 | 5 | 5 |

Finally, because these error effects occurred for a subset of the signs (two-handed signs with hand contact), we analysed response latencies only for these signs. If SPL is engaged in overt motor activity, but only for signs with the additional demands related to a body-anchored place of articulation, the above analysis including all signs may have missed such effects. This 2×2 ANOVA revealed no main effect of coil location (F_1_ < 1, F_2_ (1,59)=1.210, p=.276), and the main effect of TMS was significant only by items (F_1_(1,13) = 4.286, p=.059; F_2_(1,59) = 4.160, p=.046): a non-reliable tendency for faster responses to TMS trials regardless of stimulation location. Crucially there was no significant interaction (both F < 1).

## 4. Discussion

TMS to SPL had very specific effects: an increased rate of phonological feature substitution errors for complex two-handed signs (those requiring hand contact), but did not slow or otherwise impair performance. The lack of effects of TMS on response initiation times suggests that SPL is not engaged in execution, but may instead be engaged in generation of internal motor plans that are used for error correction/prevention as hypothesized by Emmorey et al. (2004; 2007; 2014; 2016). TMS also did not affect the rate of dysfluencies, in contrast to studies involving reaching and grasping which indicate a finer-grained role of SPL in motor control. Our results support the hypothesis that SPL is particularly linked to body-centric motor representations and is engaged when movement must reach a particular target location.

The specific pattern of results allows us to draw further conclusions about the role of SPL and self-monitoring in sign production: TMS to SPL decreased the likelihood of detecting or correcting phonological errors during otherwise successful bimanual coordination, but it did not affect errors of other types nor other kinds of signs. The lack of TMS effects on dysfluent productions (i.e., slight variation in fine details of hand configuration, movement or place of articulation) may come as a surprise, given the various results from studies of reaching and grasping (Jackson and Husain, 2006; Parkinson et al., 2010; Reichenbach et al., 2014) in which SPL seems to be clearly involved in fine motor control and correction of perturbations. There are, however, a number of reasons why this may not have been the case in the present study. First, as lexical signs are made up of combinations of contrastive discrete elements (hand configuration, movement, place of articulation), producers have considerable leeway in the precise formation of any given sign (Brentari, 1998; Corina and Sandler, 1993) which is reflected in articulatory variation between tokens of the same sign as observed in corpora of natural signing (Fenlon et al., 2013). Such variation is especially prevalent when signs are produced in sentence contexts, where coarticulation effects also have substantial consequences on the fine-grained motor details of sign form (Corina and Sandler, 1993). Second, for similar reasons, fine motoric deviations from a signer’s intended production may not be noticeable by a comprehender. Such fine deviations may have existed in our dataset, but went unnoticed by the coder because they fell within the “acceptable” range of possible variation in producing a given sign (c.f. false positives in tasks involving nonsense signs, Orfanidou et al., 2015), while phonological feature errors were simply more evident. Finally, the lack of visual guidance in sign language production may fundamentally change the nature of the motor representations that are needed for correct production, compared to other motor tasks where vision plays an essential role. It may be tempting to conclude that for sign language production, for which visual guidance is not necessary and motor plans are retrieved from memory rather than being determined by the physical environment, the role of SPL has shifted to generation of phonologically-based forward models at the expense of finer control. However, in the present study (as well as Emmorey et al., 2016) SPL activity is prevalent for a subset of signs: those involving contact between the hand and body. These are cases where finer control is necessary to reach a target location, although some variation is still permitted. We suggest that SPL is carrying out a dual motoric role, both generation of fine-grained forward models of motor action, as well as generation of phonemic forward models. For signs that have increased demands on fine-grained aspects of movement (in particular, reaching a more precisely defined final state in contact with the body, compared to signs without such contact), phonological monitoring may increase the processing burden, and thus may be more susceptible to interference due to TMS. As the finer-grained motor components are resilient to minor perturbation in the case of sign production, a corresponding rise in dysfluencies may not occur (or may not be detectable). Ultimately, for fluent signers SPL has adapted to monitor motor plans for discrete hand configurations retrieved from memory as well as more fine-grained aspects of visually guided actions.

---

**Supplemental Table 1.**
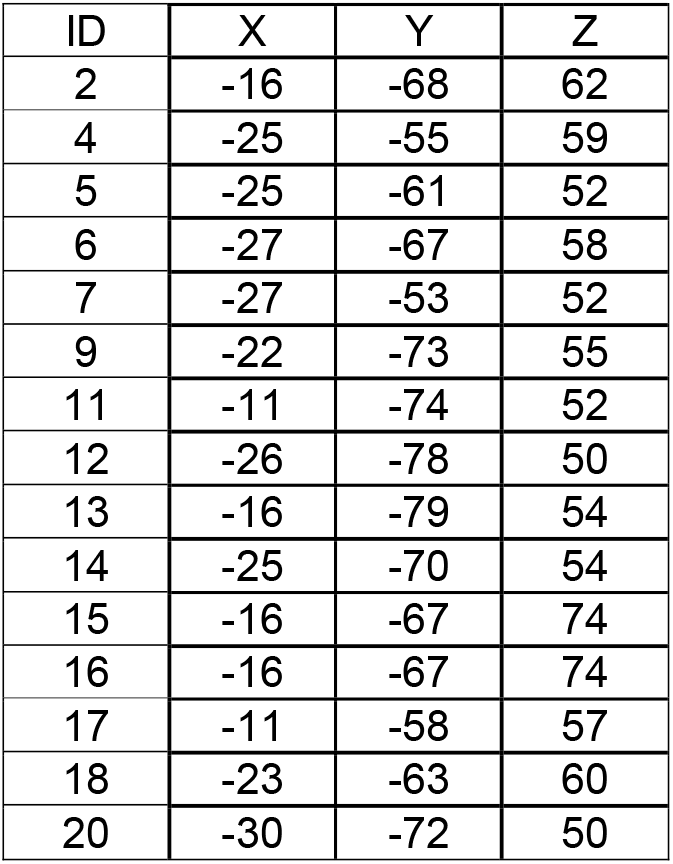
Coordinates (MNI) of peak activity for covert naming > rest in left SPL for individual participants in preliminary fMRI study. These coordinates were targeted in the TMS study.

Supplemental Data. Trial-level data from the BSL naming study including response times and error classification (CSV format). Subject: participant identifier. NamingLatency: Key release time (milliseconds). NamingLatency2: Key release time (millseconds), second hand released (not used in analyses). Target: label of target picture. CoilLocation: position of TMS coil (S: SPL, V: Vertex). TMS: Presence of TMS (0/1). Accuracy: Correct or incorrect (0/1); NA indicates no sign was produced. Dysfluency: Dysfluent production (0/1). Phonological: Phonological feature error (0/1). Lexical: Lexical error (0/1). Handed: Sign produced with 1 or 2 hands (can vary between individuals for the same concept). Contact: Indicates whether the sign involves hand contact, body contact or no contact. Handshape: For two-handed signs, indicates whether the handshapes are same or different. Movement: For two-handed signs, indicates whether the hand movement is the same for both hands, same but alternating, or involves only movement of the dominant hand. ProducedSign: Indicates whether a sign was produced on that trial.

